# Identifying and validating housekeeping hybrid *Prunus* sp. genes for root gene-expression studies

**DOI:** 10.1101/611004

**Authors:** Adriana Bastías, Kristen Oviedo, Rubén Almada, Francisco Correa, Boris Sagredo

**Author notes:** Corresponding author: Boris Sagredo, Instituto de Investigaciones Agropecuarias (INIA) CRI Rayentué, Av. Salamanca s/n, Sector Los Choapinos, Rengo, Chile.

## Abstract

Prunus rootstock belonging to subgenera *Amygdalus* (peach), *Prunus* (plum) and *Cerasus* (cherry) are either from the same species as the scion or another one. The number of inter-species (including inter-subgenera) hybrids have increased as a result of efforts to broaden the genetic basis for biotic and abiotic resistance/tolerance. Identifying genes associated with important traits and responses requires expression analysis. Relative quantification is the simplest and most popular alternative, which requires reference genes (housekeeping) to normalize RT-qPCR data. However, there is a scarcity of validated housekeeping genes for hybrid Prunus rootstock species. This research aims to increase the number of housekeeping genes suitable for Prunus rootstock expression analysis.

Twenty-one candidate housekeeping genes were pre-selected from previous RNAseq data that compared the response of root transcriptomes of two rootstocks subgenera to hypoxia treatment, ‘Mariana 2624’ (*P. cerasifera* Ehrh.× *P. munsoniana* W. Wight & Hedrick), and ‘Mazzard F12/1’ (*P. avium* L.). Representing groups of low, intermediate or high levels of expression, the genes were assayed by RT-qPCR at 72 hours of hypoxia treatment and analyzed with NormFinder software. A sub-set of seven housekeeping genes that presented the highest level of stability were selected, two with low levels of expression (*Unknown 3, Unknown 7*) and five with medium levels (*GTB 1, TUA 3, ATPase P, PRT 6, RP II*). The stability of these genes was evaluated under different stress conditions, cold and heat with the hybrid ‘Mariana 2624’ and N nutrition with the hybrids ‘Colt’ (*P. avium* × *P. pseudocerasus* Lindl.) and ‘Garnem’ [*P. dulcis* Mill.× (*P. persica* L.× *P. davidiana* Carr.)]. The algorithms of geNorm and BestKeeper software also were used to analyze the performance of these genes as housekeepers.

Stability rankings varied according to treatments, genotypes and the software for evaluation, but the gene *GBT 1* often had the highest ranking. However, most of the genes are suitable depending on the stressor and/or genotype to be evaluated. No optimal number of reference genes could be determined with geNorm software when all conditions and genotypes were considered. These results strongly suggest that relative RT-qPCR should be analyzed separately with their respective best housekeeper according to the treatment and/or genotypes in *Prunus* spp rootstocks.

## Introduction

Stone fruit trees (*Prunus* sp.) are economically important because they produce edible fruits such as peaches, cherries, and apricots. Most stone fruit trees are grafted on rootstocks (seedlings or clonally propagated) that belong to either the same or other *Prunus* species [1]. Therefore, these fruit trees are composed of two parts, the rootstock and scion. Rootstocks are responsible for absorbing water and nutrients and providing resistance to soil pathogens, pests and tolerance to environmental conditions. Adaptation to environmental stressors like drought, salinity, cold, and root hypoxia are largely determined by the rootstock [2]. Other important attributes of fruit cultivars like initial flowering, vigor, nutritional state, fruit production, size, and taste can be significantly influenced by the rootstock [3,4]. To be grafted, rootstocks must be compatible with a wide range of cultivars, resistant to pests and diseases, and suited to different soil types [5]. It is unlikely that any single Prunus genotype has all these attributes. Incorporating the maximum number of these characteristics to increase usefulness and areas of adaptation is the main goal of Prunus rootstock breeding programs [6]. However, there are important limitations to traditional breeding programs, including long generation times and extensive space requirements. Deepening our understanding of the molecular basis of the traits/responses of Prunus rootstocks and identifying the genes involved will be critical steps toward improving rootstock plants by marker-assisted selection (MAS) methods [7]. Other approaches are also possible, based on direct modification of key genes associated with traits of interest by genetically engineering Prunus species [8]. However, the molecular basis of Prunus rootstock traits and responses remain largely unknown. Gene expression studies using NGS (Next Generation Sequencing) and/or candidate genes from other model plants represent valuable approaches to identifying key candidate genes underlying traits of interest [9].

Quantitative real-time PCR (RT-qPCR) is a powerful tool for quantifying gene expression due to its high degree of sensitivity, specificity and reproducibility [10]. Normalizing RT-qPCR data is crucial to obtain results as close to reality as possible. Biological (gene-specific) and technical (RNA quantity and quality, RT efficiency and PCR efficiency) variations occur in gene expression analysis. Appropriate normalization strategies are required to control experimental error during the multistage process, which also include extracting and processing RNA samples [11]. There are several qPCR methods of absolute and relative quantification. Absolute quantification, which require normalizing to sample size or a standard curve, quantifies transcript in a given sample [12]. Relative quantification analyzes changes in gene expression in a given sample relative to a reference sample. Among several proposed methods, reference genes are frequently used to normalize RT-qPCR data [11,12]. Correct normalization of gene expression levels requires identifying housekeeping, reference or control genes, the expression levels of which are unaffected by experimental factors [11]. The expression of housekeeping genes should vary minimally among different tissue and physiological states of the organism [13]. Housekeeping genes function as internal controls that are subject to the same conditions as the mRNAs of interest, which are also measured by real time RT-qPCR [11]. The simplest and most popular method is relative quantification, which requires reference or housekeeping genes to normalize RT-qPCR data [10]. A serious limitation of this type of gene expression study of Prunus rootstocks is the scarcity of validated reference genes. Because modern Prunus rootstocks include genotypes of different peach, plum and cherry subgenera, including several inter-specific hybrids, identifying reference genes with stable expression among *Prunus* sp. species is a challenge.

Several plant genes have been used as internal controls in expression studies, such as glyceraldehyde-3-phosphate dehydrogenase (*GAPDH*), tubulin (*TUB*), actin (*ACT*) and 18S ribosomal RNA (*18S rRNA*) [13,14]. However, there are no universal reference genes with constant levels of expression for all plants, tissue, treatments and developmental stages. Stable expression in one organism or species may not be suitable for standardization in another, or for the same tissue but under different conditions [13]. Hence, researcher must determine the best housekeeping genes according to the specific experimental conditions. Identifying these genes is not a simple process. It consists of two steps: first, identifying likely candidate genes and then determining their stability [15].

In this work, we searched for and validated housekeeping genes for Prunus rootstock gene expression analysis. We assessed candidate housekeeping genes from our existing RNAseq data [16] and in the literature [17]. The qRT-PCR data were analyzed using three widely applied algorithms NormFinder, geNorm and BestKeeper, to determine suitable *Prunus sp* housekeeping genes for six experimental conditions: hypoxia, drought, salinity, cold, heat and N nutrition.

## Materials and Methods

### Plant material

Clonally propagated and virus-free rootstock plants from Mariana 2624 (*P. cerasifera* × *P. munsoniana* W. Wight & Hedrick), Mazzard F12/1 (*P. avium*), Colt (*P. avium* (L.) L. × *P. pseudocerasus* Lindl.) and Garnem [*P. dulcis* × (*P. persica* × *P. davidiana*)] were donated by a commercial nursery (Agromillora Sur, S.A., Curicó, Chile). Plants were transplanted to 2-L plastic pots with a mixture of vermiculite:perlite:sand (1:1:1v/v) as a substrate. The plants were maintained in the field under a shade net (Raschel sun shading net with 50 % shading) at the Instituto de Investigaciones Agropecuarias - Rayentué (Rengo, Chile) during two growing seasons until they were used in different experiments. The plants were watered three times a week with tap water and fertilized every 2 weeks with 1 g/pot with N:P:K (25:10:10) (Ultrasol™, Soquimich, Chile).

### Abiotic stress treatments

Hypoxia experiments as described by Arismendi et al. [16] were carried out. With the exception of the control plants, the plants in their pots were placed in 100-L plastic containers when they reached an average height of 30 cm. Root hypoxia was generated by filling the plastic containers with water until approximately 4 cm above the level of the pot substrate, and then bubbling 100 % gaseous N2 (1 L/min) through the water to displace dissolved O2. The oxygen levels in close proximity to the plant roots were monitored throughout the experiment with an oxygen-electrode (Extech Instrument, MA, USA). With a total of 15 plants per genotype, roots from three randomly selected plants were collected at 0 and 72 h. Samples collected at time 0 h represented the control without flooding. To take root samplings, the soil was completely removed and the plant roots were gently washed with tap water, then excised from the plants, immediately frozen in liquid nitrogen, and stored at −80 °C until RNA extraction.

Saline, cold, heat and drought stress treatments were applied according to [18,19]. ‘Mariana 2624’ plants of uniform size (30 cm tall) were used for these experiments. The roots of ten plants were immersed in a 200 mM NaCl solution for 0, 6 and 24 h for the saline treatment. For the cold and heat treatments, plants were kept in growth chambers at respectively 4 and 37 °C for 0, 6 and 24 h. Finally, for the drought treatment, plant roots were washed gently with water to remove soil and then put in perlite for rapid dehydration [18]. Roots from three plants were collected from the control (without treatment) and treated plants at 6 and 24 h. The removed roots were immediately frozen in liquid nitrogen and stored at −80 °C for RNA extraction and gene expression analysis.

### Nitrogen (N) Treatment

‘Garnem’[*P. dulcis* × (*P. persica* × *P. davidiana*)] and ‘Colt’ (*P. avium* × *P. pseudocerasus*) plants were transplanted to 2-L plastic pots with a mixture of vermiculite:perlite (1:1 v/v) as substrate. Plants were maintained in the field under a shade net (Raschel sun shading net with 50 shading) at the Instituto de Investigaciones Agropecuarias - Rayentué (Rengo, Chile) during two consecutive growing seasons (2013-2014 and 2014-2015). Plants were watered twice a week, once with tap water and the other with a modified Murashige & Skoog nitrogen-free basal medium (Phytotechnology Laboratories, M531), supplemented with 0.0, 1.0 and 10.0 mM of ammonium nitrate (NH4NO3) for two consecutive months of plant growth during two seasons. The roots of three randomly selected plants of each genotype and treatment with different N doses were sampled at the end of both seasons. Control plants were maintained under standard irrigation conditions, with watering twice a week with tap water, and fertilizing every two weeks with 1 g/pot of N:P:K (25:10:10) (Ultrasol™, Soquimich, Chile). For root samplings, the soil was completely removed and plant roots were gently washed with tap water, excised from the plants, immediately frozen in liquid nitrogen and stored at −80 °C until RNA extraction.

### Gene expression analysis by RT-qPCR

Total RNA of one plant per treatment and three control plants was used at sampling times for gene expression analysis by quantitative PCR (RT-qPCR). The RNA extraction procedure was as described by Arismendi et al. [16]. Total RNA was extracted from root samples of control and treated plants according to [20]. Following the DNase treatment, 5 μg of total RNA was used to synthesize cDNA from each sample, using a Thermoscript RT-PCR System™ (Invitrogen, Inc., Carlsbad, CA). Gene transcript levels were measured by qRT-PCR using a Mx3000P QPCR System (Agilent Technologies, Santa Clara, CA). All reactions were made with the Brilliant SYBR Green Master Mix (Stratagene Inc., Santa Clara, CA), according to the manufacturer’s instructions. All qRT-PCR reactions were done in triplicate (technical replicates) using 2 μL Master Mix, 0.5 μL 250 nM of each primer, 1 μL of diluted cDNA and nuclease-free water to a final volume of 20 μL. Controls (with no cDNA and RNA without RT) were included in all runs. Fluorescence was measured at the end of the amplification cycles (Ct). Amplification was followed by melting curve analysis with continual fluorescence data acquisition from 65 to 95 °C.

### Selection of Prunus putative housekeeping genes

Twenty-one stable candidate genes were selected from a previous analysis of RNA sequencing of Prunus rootstocks [16] to be evaluated as expressed reference genes in RT-qPCR studies. Several classical reference genes described by Tong et al. [17] were also included. The previous RNA sequencing data consisted of root transcriptomes from two rootstocks genotypes, ‘Mariana 2426’ and ‘Mazzard F12/1’, with 0, 6, 24 and 72 h of hypoxia treatment [16]. Briefly, gene abundance was estimated by FPKM count (fragments per kilobase of transcript per million mapped reads) [21,22]. As a cutoff, we only considered genes with a minimum of 10 aligned fragments (-c option). To consider a gene as a candidate housekeeper, the calculated fold change among all time points had to be between −0.3 to +0.3. The coefficient variance (cv) values of less than 0.3 and FDR-corrected P values <0.05 were used as filters. Categories of levels of expression were defined to ensure representability (low (reads < 200), intermediate (200< reads < 3000) and high levels of expression (reads > 3,000). Primers for genes suitable for qPCR were designed with Primer Premier 6.0 software, with a melting temperature between 58–61°C, 21–23 bp and approximately 50% GC content. Amplicon lengths were between 160–280 bp. All primers were synthetized at IDT (Integrated DNA Technologies, Inc., CA)

### Determining stability of housekeeping gene expression and statistical analysis

The expression levels of the candidate reference genes were determined by the number of amplification cycles (Cq) needed to reach a specific threshold level of detection. The obtained data were analyzed using NormFinder [15], geNorm [23] and BestKeeper software packages [24]. RT-qPCR data were exported to an Excel datasheet, and the Ct values were converted according to software requirements. Each of these approaches generates a measure of housekeeping gene stability, which can be used to rank genes as candidate housekeeping genes according to their stability.

## Results

### Identification of putative housekeeping genes for *Prunus sp*

A total of 21 genes were selected for this study to identify housekeeping genes with the highest levels of expression for RT-qPCR studies of *Prunus sp*. Previous data analysis of RNA sequencing of root transcriptomes of two Prunus sp. rootstocks assayed under hypoxia [16] were used to select putative reference genes for Prunus sp. root expression. The genotypes ‘Mariana 2624’ and ‘Mazzard F12/1’, from *Prunus* and *Cerasus* subgenera, are respectively tolerant and sensitive to hypoxia. Using the same probabilistic model [21,22] that Arismendi et al. [16] used, the most stable genes among treatments and genotypes were identified. The analysis generated 611 candidate housekeeping genes, with relatively constant expression levels, with fold changes between −0.3 and +0.3 and CV <0.3 (Table S1). Twenty-one genes were selected to be evaluated by RT-qPCR, with 4, 10 and 7 of them respectively representing low (reads<200), intermediate (200>reads>3,000) and high levels expression (reads>3,000) (Table 1). This range of gene expression was chosen to identify suitable reference genes for better relative quantification of genes of interest with low, intermediate and high levels of expression.

**Table 1.** Description of candidate housekeeping genes and their expression level according Arismendi et al. (2015).

| **Name** | ***P. persica* database**<br>**accession number** | **RNAseq**<br>**Expression level** | **Arabidopsis**<br>**homolog locus** | ***A. thaliana* locus description** | **Protein**<br>**similarity (%)** |
| --- | --- | --- | --- | --- | --- |
| *Unknown 2* | ppa013569 | Low | unknown | unknown | - |
| *Unknown 3* | ppb022375 | Low | unknown | unknown | - |
| *Got 1* | ppa013371 | Low | AT5G24170 | Got1/Sft2-like vesicle transport protein family | 72.8 |
| *Unknown 7* | ppa026944 | Low | unknown | unknown | - |
| *CYP 2* | ppa011090 | Medium | AT4G34960 | Cyclophilin-like peptidyl-prolyl cis-trans isomerase protein | 86.2 |
| *RP II* | ppa008812 | Medium | AT2G15430 | DNA-directed RNA polymerase family protein | 92.8 |
| *3B* | ppa000339 | Medium | AT5G64270 | Splicing factor. putative | 89.7 |
| *ATPase P* | ppa000424 | Medium | AT5G23630 | Phosphate deficiency response 2 | 84.8 |
| *PRT 6* | ppa000069 | Medium | AT5G02310 | Proteolysis 6 | 60.9 |
| *VPS 13* | ppa000004 | Medium | AT4G17140 | Pleckstrin homology (PH) domain-containing protein | 76.2 |
| *PHD* | ppa000413 | Medium | AT3G02890 | RING/FYVE/PHD zinc finger superfamily protein | 33.1 |
| *LBA 1* | ppa000334 | Medium | AT5G47010 | RNA helicase. putative | 82.8 |
| *GTB 1* | ppa000164 | Medium | AT1G65440 | global transcription factor group B1 | 68.3 |
| *TUA 3* | ppa005642 | Medium | AT1G50010 | tubulin alpha-2 chain | 93.6 |
| *TUB* | ppa005644 | High | AT1G20010 | tubulin beta-5 chain | 94.7 |
| *GAPDH* | ppa008250 | High | AT3G04120 | glyceraldehyde-3-phosphate dehydrogenase | 88.5 |
| *ACT 7* | ppa007211 | High | AT5G09810 | Actin 7 | 99.5 |
| *RPL 13* | ppa011512 | High | AT3G49010 | Breast basic conserved 1 | 79.7 |
| *TEF 2* | ppa001367 | High | AT1G56070 | Ribosomal protein S5/Elongation | 96.9 |
| *Unknown 8* | ppa011598 | High | AT5G50200 | nitrate transmembrane transporters | 52.5 |
| *AAT 1* | ppa005315 | High | AT5G11520 | aspartate aminotransferase 3 | 83.9 |

The putative housekeeping genes comprise both widely-used classical reference genes like *GAPDH, TUB* and *ACT* [17], and ones less commonly used like a cyclophilin-like peptidyl-prolyl cis-trans isomerase family protein-coding genes (*CYP2*), a global transcription factor of group B1 (*GTB1*) and a Got1/Sft2-like vesicle transport protein (*GOT 1*) (Table 1).

### Evaluation of candidate housekeeping genes for *Prunus sp* under hypoxia stress

The levels of expression of the candidate housekeeping genes were determined by RT-qPCR using root RNA samples from two genotypes of *Prunus sp* that were subjected to 0 and 72 h of hypoxia stress, while their stability was evaluated with NormFinder software [15]. Table 2 shows primer sequences for the studied housekeeping genes, designed with Primer Premier 6.0 software. Quantification cycle values (Cq) of the candidate genes were represented in box and whisker plots (Figure 1) that graphically show gene Cq variation, and thus give good approximations of the best candidate housekeeping genes under these conditions and for these genotypes. Table 3 shows the rankings of candidate housekeeping genes considering overall samples (genotypes and treatments). The stability values ranged between 0.006 (SE±0.002) and 0.061 (SE±0.013). Table 3 also shows rankings that considered genotypes (‘Mariana 2624’ and ‘Mazzard F12/1’) and treatments (0 and 72 h under flooding) separately. There are seven genes highlighted in bold in Table 3 that were selected for further validation analysis because they had higher stability values.

**Table 2.**
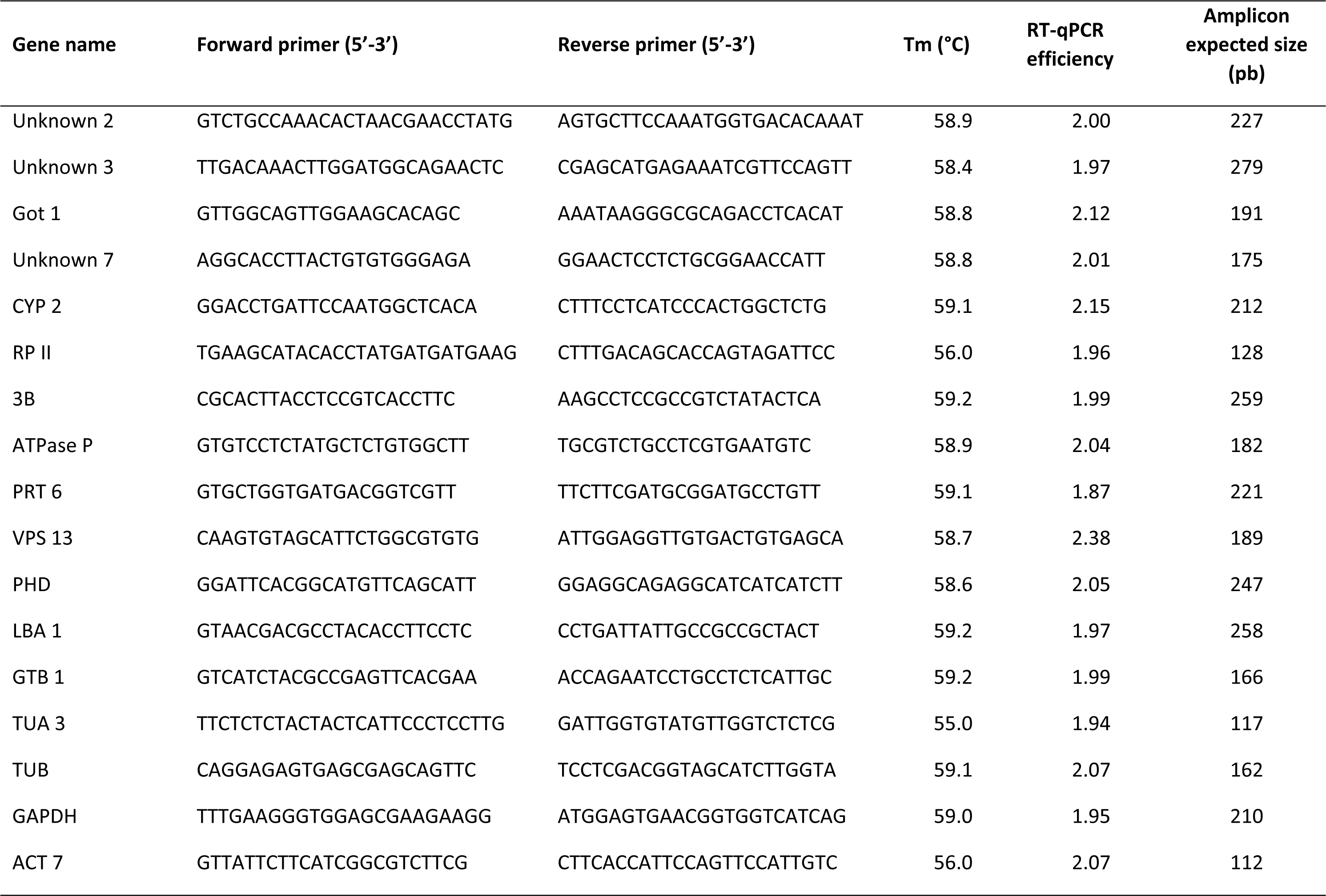
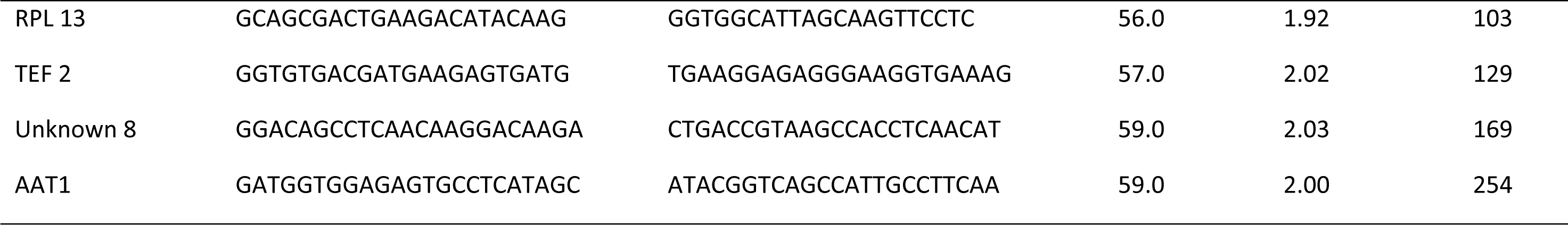
Primers sequences for each housekeeping gene studied.

**Table 3.** Ranking of putative housekeeping genes according their expression stability calculated by NormFinder software in roots of ‘Mariana 2624’ and ‘Mazzard F12/1’ under hypoxia considering genotype and treatment.

| Ranking | Genotype & Treatment | Genotype | Treatment |
| --- | --- | --- | --- |
| 1 | *RP II* | *GTB 1* | *TUA 3* |
| 2 | *GTB 1* | *Unknown 7* | *Unknown 3* |
| 3 | *TUA 3* | *ATPase P* | *PRT 6* |
| 4 | *PRT 6* | *PRT 6* | *TUB* |
| 5 | *VPS 13* | *TUA 3* | *GTB 1* |
| 6 | *Got 1* | *RP II* | *RP II* |
| 7 | *LBA 1* | *Got 1* | *RPL 13* |
| 8 | *3B* | *3B* | *VPS 13* |
| 9 | *GAPDH* | *VPS 13* | *GAPDH* |
| 10 | *RPL 13* | *RPL 13* | *LBA 1* |
| 11 | *TUB* | *AAT 1* | *Unknown 2* |
| 12 | *Unknown 7* | *Unknown 8* | *Got 1* |
| 13 | *ATPase P* | *Unknown 3* | *3B* |
| 14 | *Unknown 3* | *GAPDH* | *AAT 1* |
| 15 | *AAT 1* | *LBA 1* | *Unknown 7* |
| 16 | *CYP2* | *TUB* | *ATPase P* |
| 17 | *Unknown 8* | *ACT 7* | *CYP 2* |
| 18 | *PHD* | *CYP 2* | *TEF 2* |
| 19 | *TEF 2* | *PHD* | *PHD* |
| 20 | *ACT 7* | *TEF 2* | *Unknown 8* |
| 21 | *Unknown 2* | *Unknown 2* | *ACT 7* |

**Figure 1.**
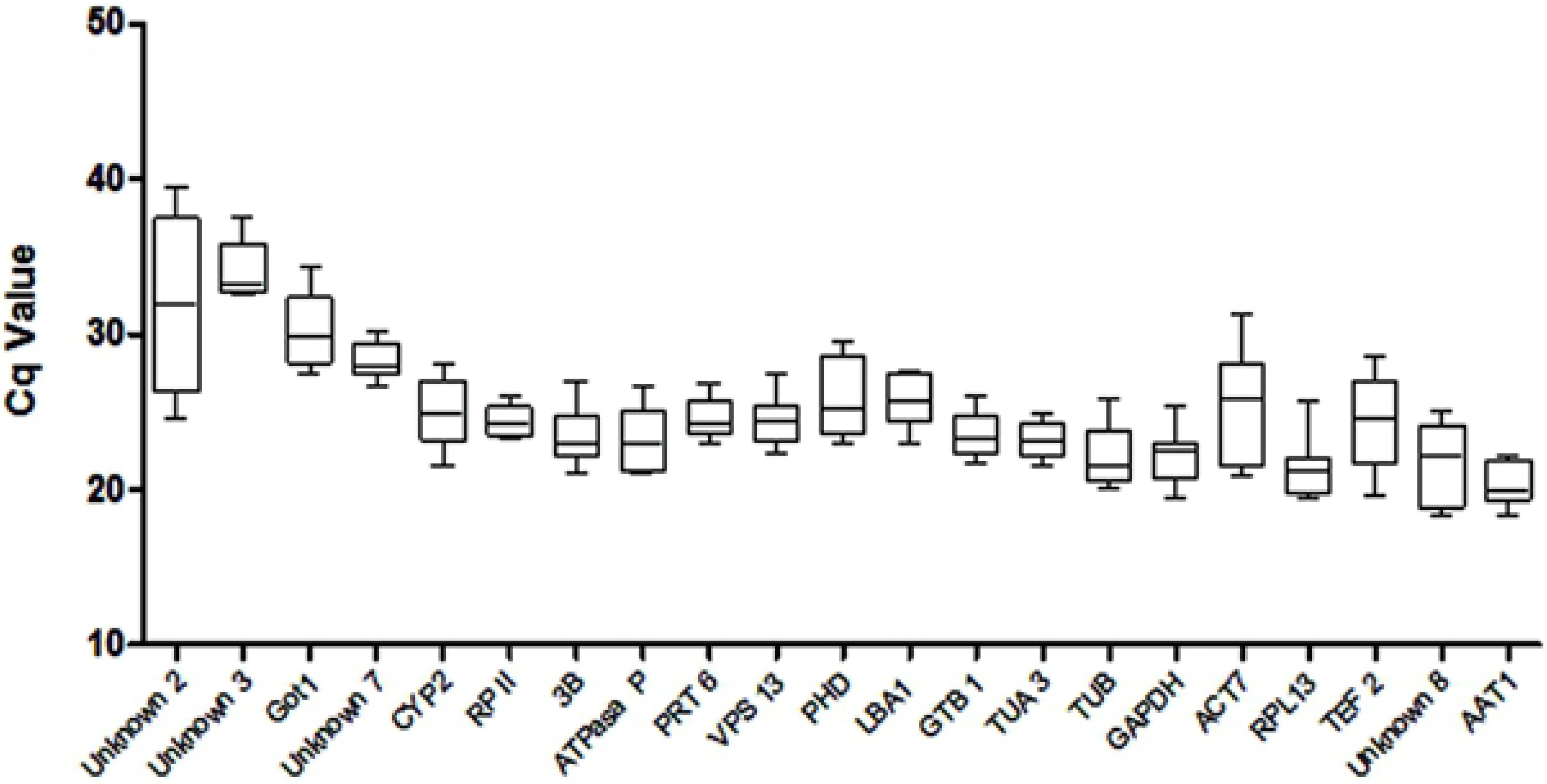
Expression levels of different candidate housekeeping genes for Prunus sp. Expression data displayed as RT -qPCR quantification cycle (Cq) values for the housekeeping genes in *Prunus* sp. under hypoxia. The line across the box depicts the median. The box indicates the 25th and 75th percentiles, and whisker caps represent the maximum and minimum values.

### Validation of candidate *Prunus sp*. housekeeping genes under new conditions

The previously selected genes, *PRT 6, GTB 1, ATPasa P, RP II, TUA 3, Unknown 3* and *Unknown 7*, were characterized to be validated as housekeeping genes under treatments with abiotic stresses, N nutrition and rootstock genotypes. The evaluation included two analytical algorithms from BestKeeper [24] and geNorm software [23]. Responses to drought, salinity, cold and heat were evaluated with the hybrid ‘Mariana 2624’ (*P. cerasifera* × *P. munsoniana*), and to N nutrition with the hybrids ‘Garnem’ [*P. dulcis* × (*P. persica* × *P. davidiana*)] and ‘Colt’ (*P. avium* × *P. pseudocerasus*), with 0.0, 1.0 and 10.0 mM of ammonium nitrate. Wang and Tsay [25] considered the first two nitrogen concentrations low and the latter high for the model plant *Arabidopsis thaliana*.

Table 4 shows the results of drought, salinity, heat and cold stress treatments with the ‘Mariana 2624’ genotype, and of the N nutrition treatment with the genotypes ‘Garnem’ and ‘Colt’, according to NormFinder software. Stability values ranged from 0.006 (*GTB 1*) to 0.064 (*TUA 3*). According to this analytical algorithm, *GTB 1* is the most stable among treatments and genotypes, followed by *Unknown 7* (Table 4). However, ATPase P was the most stable gene under the ammonium nitrate treatment, followed by the protein-coding genes *PRT 6* and *Unknown 7*.

**Table 4.**
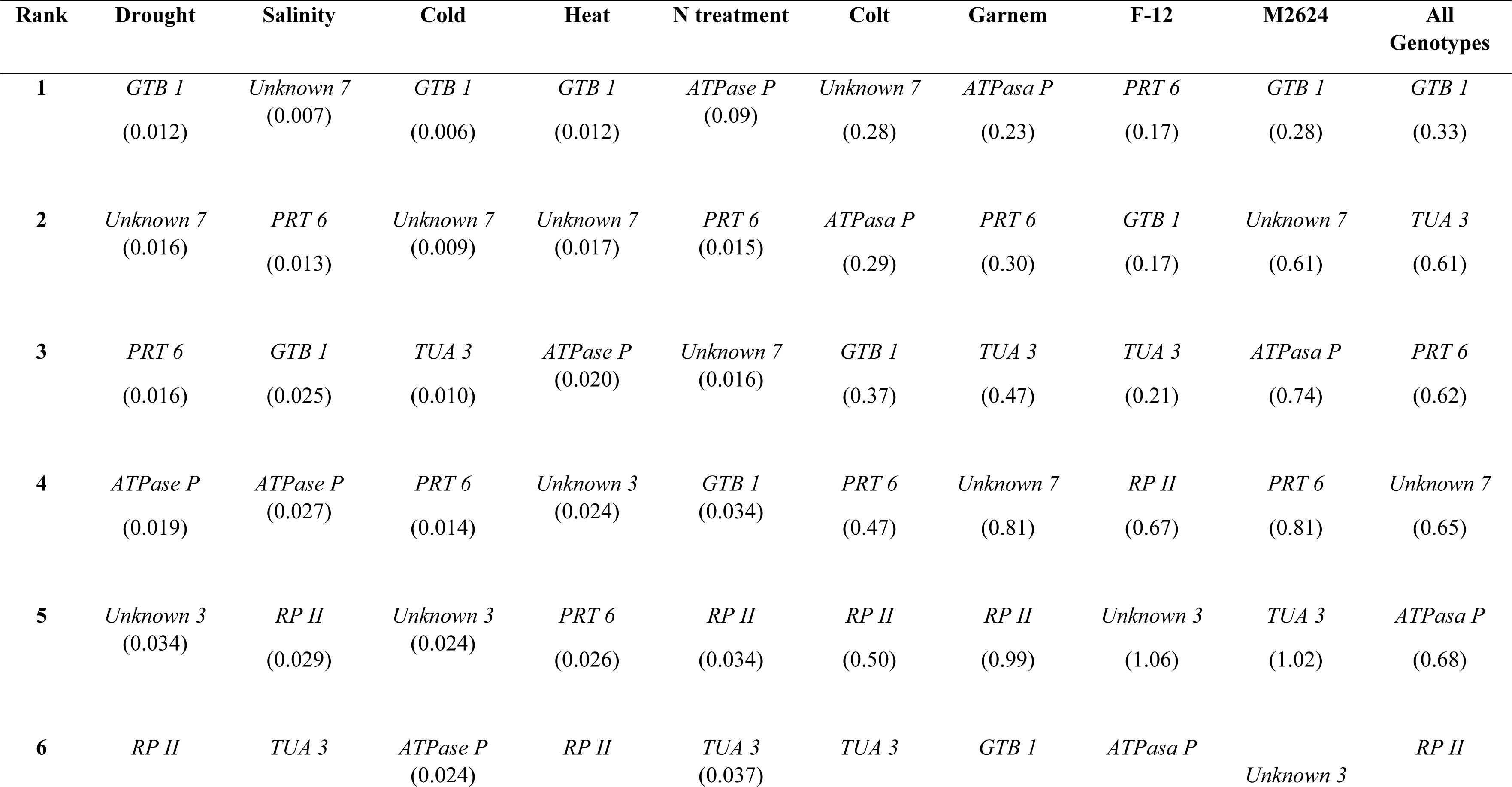
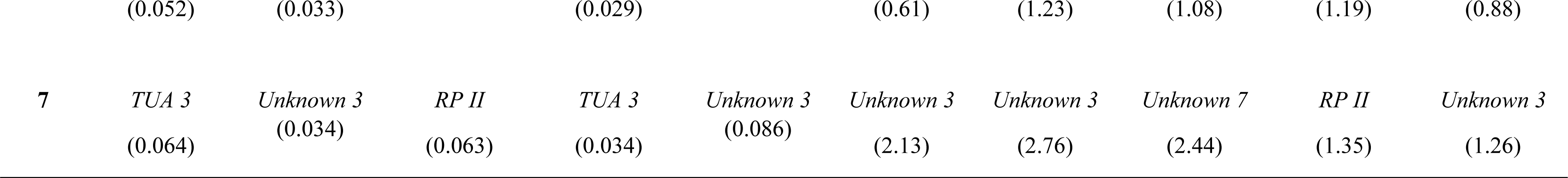
Ranking of candidate protein-coding housekeeping genes under different conditions and using distinct genotypes in order of their expression stability calculated by NormFinder software. Stability value for each condition and gene is show in brackets

The stability of the seven candidate housekeeping genes was also evaluated with BestKeeper software algorithms (Table 5) [24]. The standard deviation (SD), coefficient of variation (CV) and pair of correlation coefficients (Poisson correlation coefficient) were calculated [24]. With BestKeeper software, reference genes with an SD values greater than 1 are considered unsuitable as reference or housekeeping genes, while lower SD values indicate more stable housekeeping genes [24].

**Table 5.** Candidate housekeeping genes ranking considering each treatment and genotype. The expression stability value was calculated by BestKeeper.

| **Hypoxia** |  |  | **Colt** |  |  |
| --- | --- | --- | --- | --- | --- |
| **Rank** | **SD (± CP)** | **CV (%CP)** | **Rank** | **SD (± CP)** | **CV (%CP)** |
| *RP II* | 0.74 | 3.03 | *GTB1* | 0.66 | 2.94 |
| *TUA 3* | 0.95 | 4.11 | *Unknown 7* | 0.71 | 2.72 |
| *Unknown 7* | 1.01 | 3.57 | *RPII* | 0.71 | 2.72 |
| *PRT 6* | 1.03 | 4.18 | *ATPase P* | 0.71 | 3.1 |
| *GBT 1* | 1.03 | 4.38 | *PRT6* | 0.76 | 3.22 |
| *Unknown 3* | 1.48 | 4.36 | *TUA 3* | 0.77 | 3.36 |
| *ATPase P* | 1.90 | 8.12 | *Unknown 3* | 1.38 | 4.47 |
| **Drought** |  |  | **Garnem** |  |  |
| *Unknown 3* | 0.17 | 0.55 | *Unknown 3* | 0.16 | 0.51 |
| *PRT 6* | 0.36 | 1.61 | *PRT6* | 1.19 | 4.77 |
| *ATPase P* | 0.44 | 1.92 | *ATPase P* | 1.50 | 6.07 |
| *Unknown 7* | 0.53 | 2.00 | *TUA3* | 1.64 | 7.05 |
| *GTB 1* | 0.74 | 3.24 | *RPII* | 1.73 | 6.16 |
| *RP II* | 1.00 | 4.03 | *Unknown 7* | 1.73 | 6.16 |
| *TUA 3* | 1.03 | 4.49 | *GTB 1* | 1.97 | 8.17 |
| **Salinity** |  |  | **Mazzard F-12** |  |  |
| *Unknown 3* | 0.17 | 0.55 | *RP II* | 0.47 | 0.95 |
| *PRT 6* | 0.36 | 1.61 | *PRT6* | 0.89 | 3.67 |
| *ATPase P* | 0.44 | 1.92 | *Unknown 7* | 1.00 | 3.58 |
| *Unknown 7* | 0.53 | 2.00 | *TUA3* | 1.10 | 4.75 |
| *GTB 1* | 0.74 | 3.24 | *GTB1* | 1.12 | 4.80 |
| *RP II* | 1.00 | 4.03 | *Unknown 3* | 1.60 | 4.63 |
| *TUA 3* | 1.03 | 4.49 | *ATPase P* | 1.70 | 7.41 |
| **N treatment** |  |  | **Mariana 2624** |  |  |
| *Unknown 3* | 0.83 | 2.64 | *Unknown 3* | 0.67 | 2.10 |
| *PRT 6* | 0.98 | 4.03 | *GTB1* | 0.79 | 3.48 |
| *GTB 1* | 1.00 | 4.38 | *TUA 3* | 0.88 | 3.85 |
| *ATPase P* | 1.03 | 4.30 | *ATPase 3* | 0.88 | 3.86 |
| *TUA 3* | 1.15 | 4.97 | *Unknown 7* | 0.91 | 3.41 |
| *RP II* | 1.20 | 4.45 | *PRT 6* | 0.98 | 4.36 |
| *Unknown 7* | 1.20 | 4.45 | *RP II* | 1.05 | 4.20 |
| **Cold** |  |  | **Overall** |  |  |
| *Unknown 3* | 0.11 | 0.36 | *Unknown 3* | 0.85 | 2.65 |
| *PRT 6* | 0.19 | 0.86 | *GTB1* | 0.95 | 4.16 |
| *ATPase P* | 0.30 | 1.34 | *Unknown 7* | 1.01 | 3.78 |
| *GTB 1* | 0.32 | 1.46 | *TUA 3* | 1.01 | 4.43 |
| *TUA 3* | 0.44 | 2.03 | *ATPase P* | 1.02 | 4.42 |
| *Unknown 7* | 0.52 | 1.96 | *PRT 6* | 1.24 | 5.36 |
| *RP II* | 1.53 | 6.07 | *RP II* | 1.43 | 5.66 |
| **Heat** | | | Abbreviations: SD [± CP], standard deviation of the CP; CV [%CP], coefficient of variance expressed as a percentage on the CP level; CP, crossing point values. | | |
| *Unknown 3* | 0.16 | 0.50 |  |  |  |
| *GTB 1* | 0.43 | 1.95 |  |  |  |
| *ATPase P* | 0.52 | 2.34 |  |  |  |
| *PRT 6* | 0.54 | 2.48 |  |  |  |
| *Unknown 7* | 0.55 | 2.13 |  |  |  |
| *RP II* | 0.60 | 2.52 |  |  |  |
| *TUA 3* | 0.62 | 2.75 |  |  |  |

The best two candidate housekeeping genes under hypoxia were *RP II* and *TUA 3*, while the best two under drought, salinity, cold and N nutrition conditions were *Unknown 3* and *PRT 6*. Finally, the best two under heat were *Unknown 3* and *GTB 1*.

The best housekeeping genes for the genotypes were analyzed with Norm Finder and BestKeeper software (Tables 4–5). Norm Finder software identified different housekeeping genes as the most suitable for each genotype. *GTB 1* was the most suitable gene for the ‘Mariana 2624’ genotype, including the analysis that considered all the genotypes (Table 4). Best Keeper software identified *Unknown 3* as the most suitable for the ‘Mariana 2624’ and “Garnem” genotypes, and as first when the analysis included all the genotypes: ‘Mariana 2624’, ‘Mazzard F12/1’, “Colt” and “Garnem”.

The consistency of expression of seven candidate housekeeping genes was evaluated with geNorm software [23]. The stress conditions were analyzed separately and then overall treatment conditions and genotypes were analyzed together. Figure 2 shows the results of this analysis. The best candidate housekeeping genes are those with expression M-values close to zero. The reference parameters are applicable when the threshold M-value is lower than 1.5 [23]. Most candidate genes had stability M-values lower than 1.5, indicating their suitability as housekeeping genes under these conditions. According to geNorm software, two housekeeping or reference genes is optimal under hypoxia, drought, cold, heat and N nutrition conditions (Figure 3). An optimal normalization factor can be calculated as the geometric mean of the best housekeeping gene according to the studied condition (Figure 2). However, four housekeeping genes is optimal under salinity 4 (Figure 3). An optimal normalization factor can be calculated as the geometric mean of the housekeeping genes: *RP II, TUA 3, GBT 1* and *Unknown 7*.

**Figure 2.**
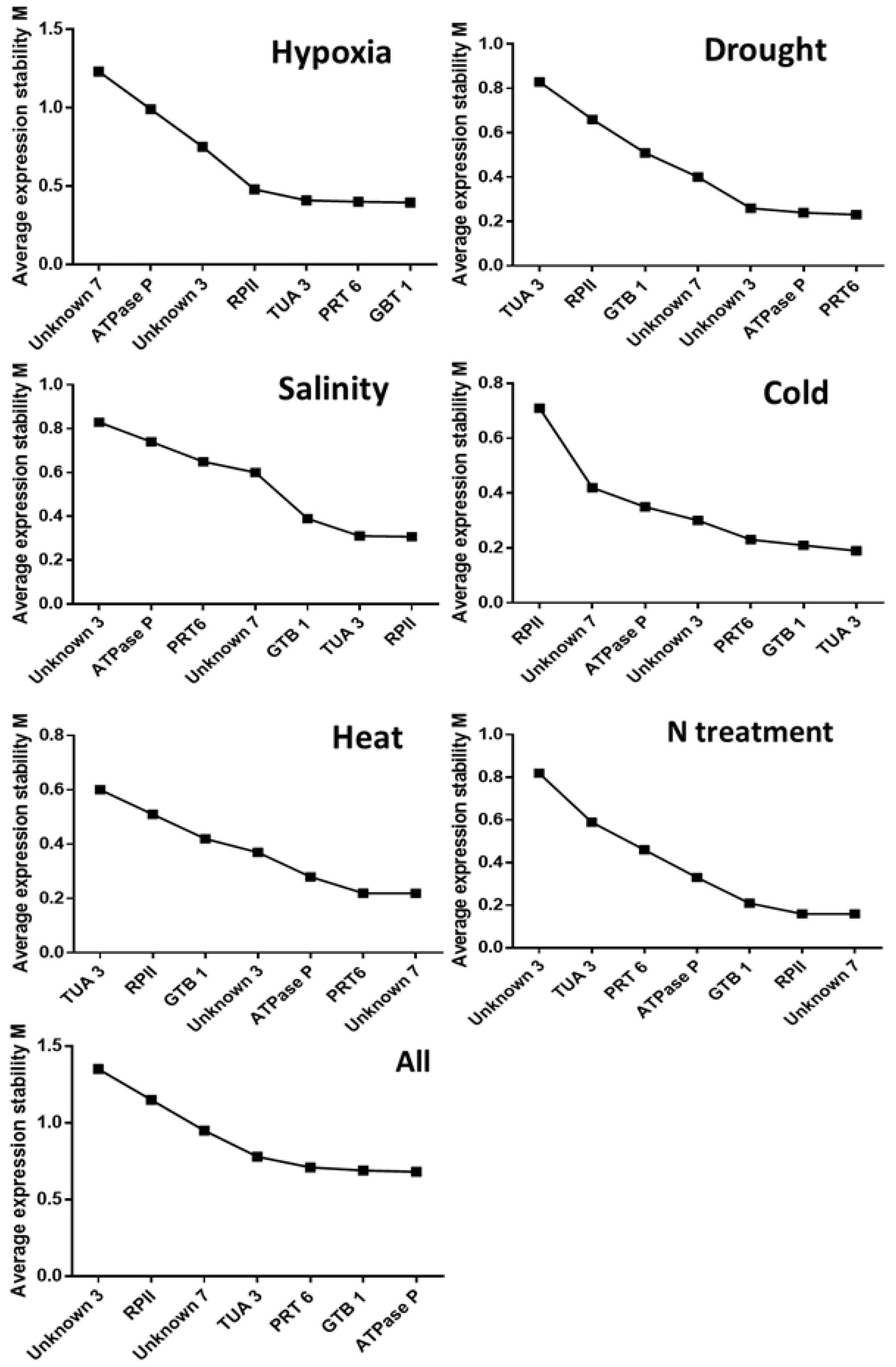
Average expression stability (M value) of the seven selected protein-coding candidate housekeeping genes using geNorm software. Expression stability was evaluated in samples from *Prunus* spp. under drought, salinity, cold, heat, N treatment and all together, plus the results of hypoxia. A lower average expression stability M value indicates more stable expression.

**Figure 3.**
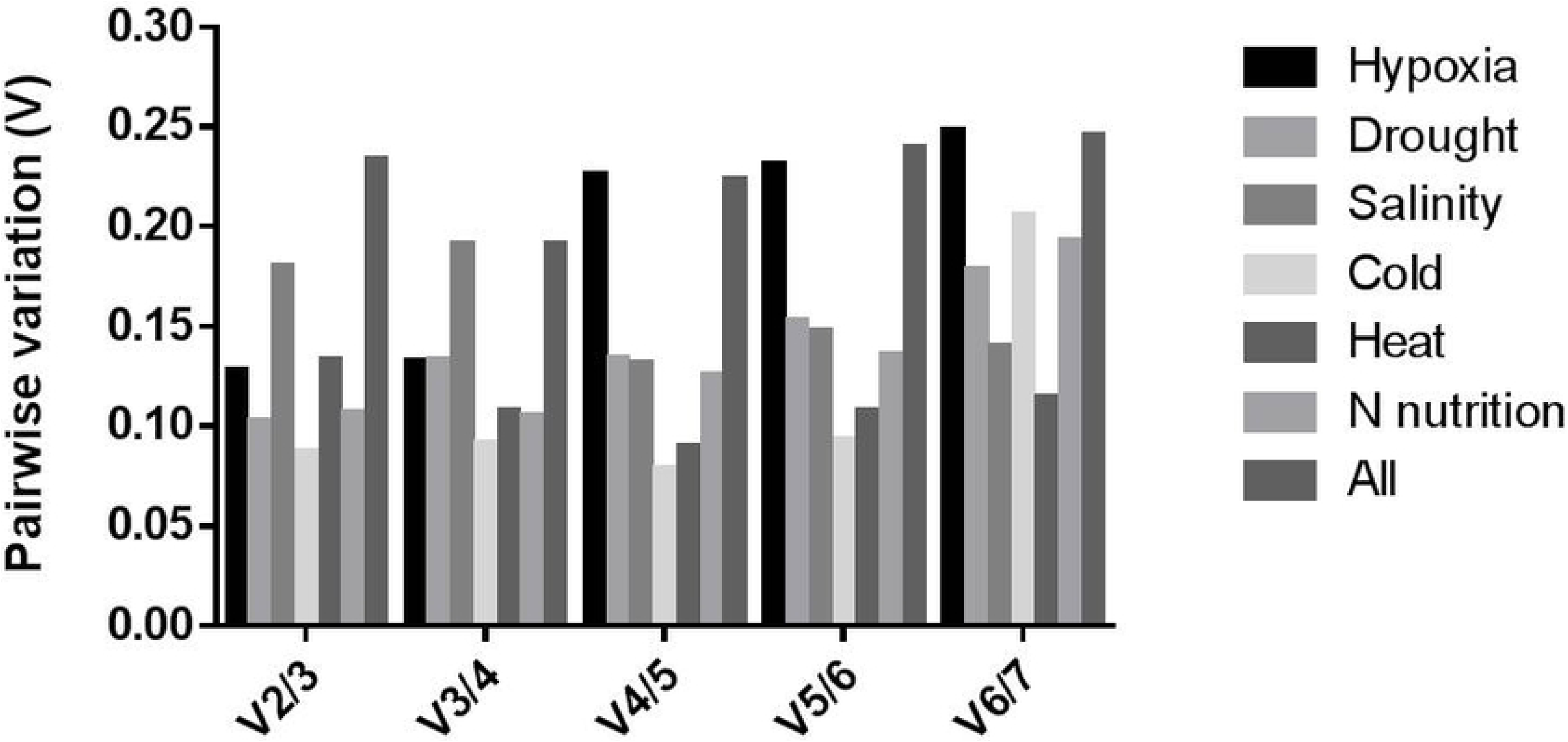
Pairwise variation (V) analysis of the seven candidate housekeeping genes using geNorm software. The pairwise variation (Vn/Vn+1) to determine the optimal number of housekeeping genes required for RT-qPCR data normalization. The cutoff value is 0.15.

Finally, Tables 4 and 5 and Figure 2 show rankings with all the Cq data obtained with NormFinder, BestKeeper and geNorm. These results include those of the seven selected housekeeping genes considering all treatments and genotypes. NormFinder and geNorm software identified *GBT 1* as the best candidate gene when all conditions and genotypes were evaluated (Table 4, Figure 2), while BestKeeper identified *GBT 1* as the second-best candidate gene (Table 5).

The cut-off value to determine the optimal number of reference genes for qRT-PCR normalization using geNorm software is 0.15. A value under 0.15 indicates that additional reference genes are not required [23]. The optimal number of reference genes could not be determined with geNorm when all conditions and genotypes were considered. This strongly suggests that relative RT-qPCR should be analyzed separately with the respective best housekeeper genes according to the treatment and/or *Prunus* spp rootstock genotype.

## Discussion

Prunus rootstocks confer adaptation to specific soil conditions, so important traits of fruit trees related to environmental stressors, such as drought, salinity, cold, and root hypoxia, are strongly and/or partially influenced by rootstock. The rootstock is often from the same subgenus, and even the same species, as the scion, *Amygdalus* (peaches), *Prunus* (plums) and *Cerasus* (cherry). Inter-species and inter-genus hybrids are increasingly common as the result of efforts to broaden the genetic basis of breeding programs to improve biotic and abiotic resistance/tolerance. Knowledge about the genetic and molecular bases of important rootstock traits is highly desirable in order to reduce breeding periods that can easily exceed 20 years. Identifying and validating genes associated with important traits and responses require knowledge of their expression pattern under different environmental conditions and treatments. The simplest and most popular method to asses gene expression is relative quantification, which requires reference genes to normalize RT-qPCR data. However, the scarcity of validated housekeeping genes for hybrid Prunus rootstock species represents a serious limitation.

A housekeeping gene is an internal standard that is assumed to remain constant among experimental groups [26]. A significant error in estimating the expression of the chosen housekeeping genes increases noise in the assay and makes it impossible to detect small changes. Worse yet, if the expression of the housekeeping gene is altered by the experimental conditions under study, the results may be entirely incorrect. Therefore, it is essential to validate potential housekeeping genes to establish whether they are appropriate for a specific experimental purpose [26].

Studies have shown that the expression patterns of many classical housekeeping genes, such as *Actin, β-TUB, GAPDH*, and *eEF-1a*, can vary under certain conditions [27,28]. Microarray and transcriptome databases are excellent resources to search for new candidate housekeeping genes, and have made it possible to identify several genes with more stable expression than classical reference genes [28,29]. Searching for housekeeping genes in microarray and transcriptome databases consists of identifying expressed genes that remain constant in the context of different test conditions, tissue, genotypes, and other factors [30]. In the present study, 611 candidate housekeeping genes were identified from RNAseq data from our previous work that compared root transcriptomes of two rootstocks ‘Mariana 2426’ (*P. cerasifera* × *P. munsoniana*) and ‘Mazzard F12/1’ (*P. avium*) under hypoxia treatments. They were filtered with fold-change expression levels between −0.3 and +0.3 and CV <0.3 among all the treatments (Table S1). Zhou et al. (2017) used more stringent criteria to search for housekeeping genes from a RNA-seq data set of apple rootstock, namely fold change values of −0.1 to +0.1, with the same CV <0.3. We identified only 15 genes as housekeeping candidates (data not shown) using this criteria. We used a less strict filter because our interest was to capture a broad range of expression of candidate housekeeping genes. Under our experimental conditions, relative categories of expression were defined according to the number of reads from our previous RNAseq experiment, with <200 reads as a low level of expression, 200> to 3,000 as an intermediate level, and >3,000 as a high level. Although the number of reads depends directly on the depth of the sequence experiment, the classification of candidate genes according in our study was according to their relative expression. These ranges of gene expression were chosen to find suitable reference genes for better relative quantification of genes of interest with low, intermediate or high levels of expression. Of the 21 candidate genes, 4 had low levels of expression, 10 had intermediate levels, and 7 high levels (Table 1).

Eight of the selected candidate genes are protein coders (ACT, CYP2, RPII, RPL13, GAPDH, TUB, TUA and TEF2) that were recommended by Tong et al [17], who identified reference genes to study gene expression in different tissue of *P. persica* with RT-qPCR. But these genes only represent the categories of intermediate (*CYP 2, RP II, TUA*) and high levels of expression (*TUB, GAPDH, ACT, RPL 13, TEF 2*).

The 21 protein-coding genes were analyzed with NormFinder. All candidate housekeeping genes can be ranked based on intra- and inter-group variations, which in our case were hypoxia treatments (0 and 72 h) and two genotypes (‘Mariana 2426’ and ‘Mazzard F12/1’). The results of the two were combined into stability values for the candidate genes (Table 3). The genes with lower stability values are more likely to be stable under any given condition [15]. The expression of stability is dependent on experimental conditions [31]. Seven of the genes with low stability values according to NormFinder were selected for further analysis (Table 3). Included were two genes with low levels of expression (*Unknown 3* and *Unknown 7*) and five with intermediate levels (*RP II, ATPase P, PRT 6, GTB 1*, and *TUA 3*). Only two of the seven (*RP II, TUA 3*) were recommended by Tong et al [17].

Codified proteins of Unknown 3 and Unknown 7 have functions that remain undetermined, RP II, RNA polymerase II is an enzyme responsible for catalyzing the transcription of gene-encoding proteins; ATPase P is a protein in the plasma membrane that maintains homeostatic balance; PRT 6 is a protein that regulates the destruction of other proteins that are no longer needed by the cell; GTB 1 is a protein responsible for regulating transcription elongation, and TUA 3 is a globular protein that is part of the microtubule, the main structural component of the cytoskeleton. The TUA protein allows transport through the cell, together with β-tubulin.

The expression of the seven genes was evaluated under new treatments and with new genotypes. The new experimental conditions consisted of subjecting the hybrid genotype ‘Mariana 2624’ to heat, cold, drought and salinity stress, and the hybrids ‘Garnem’ (*P. persica* x *P. davidiana*) and ‘Colt’ (*P. avium* x *P. pseudocerasus*) to different doses of ammonium nitrate.

The Cq values of the seven genes under new experimental conditions were determined with NormFinder, BestKeeeper and geNorm (Tables 4–5, Figure 2). The results of validating housekeeping genes generally varied according to the software, which was expected because they use different statistical algorithms. This was also observed by [31–33].

NormFinder measures variation in function of variance and ranks putative housekeeping genes according to how they differ among and within groups, while avoiding co-regulated reference genes. BestKeeper and geNorm both use the geometric mean, but BestKeeper uses raw data, and analyses up to 10 references genes, while geNorm determines stability M-value using the average pairwise variation of each candidate gene [15,23,24].

According to NormFinder, *GBT1* and *Unknown 7* ranked the highest under drought, cold, and all analyzed conditions, and all genotypes. The best housekeeping gene under salinity was *Unknown 7*, followed by *PRT 6* and *GBT 1*. Finally, the best two candidate protein-coding genes in the nitrogen treatment were *ATPase P* and *PRT 6*.

The genes *Unknown 3* and *PRT 6* ranked as the best housekeeping genes under drought, salinity, cold and N nutrition conditions according to BestKeeper analysis. *PRT 6* was the most often the best housekeeping gene, followed by *Unknown 3* (Table 5). *Unknown 3* was the only suitable housekeeping gene for the Garnem genotype, but was also suitable for the Mariana 2624 genotype, and when all the genotypes and treatment were considered (Table 5).

All genes analyzed by geNorm software under different experimental conditions had stability M-values <1.5, and were suitable as reference genes. This statistical tool predicts the optimal number of housekeeping genes necessary to normalize the experiment. Most treatments analyzed with geNorm software in this study need two reference genes, except under conditions of salinity, where four genes are required for normalization (Figure 3).

High- or middle-ranking genes generally vary slightly, depending on the analytical algorithm used (Tables 4–5 and Figure 2). In contrast, we observed more coincidence among the poorest performing housekeeping genes under the different conditions. For example, *TUA 3* and *RP II* performed the poorest among the seven genes under drought and heat conditions according the three analytical programs (Tables 4-8, Figure 2).

According geNorm software, no optimal number of reference genes could be determined when all the conditions and genotypes were analyzed together, because variability between sequential normalization factors (based on the n and n+1 least variable reference targets) is relatively high (geNorm V > 0.15). This result concurs with several studies that indicated that different housekeeping genes must validated in accordance with the experimental conditions (Figure 3).

We validated seven housekeeping genes in this study for use in qPCR analysis of *Prunus sp*. roots. Most of these genes had not been described before in Prunus, but two (RP II and TUA 3) were recommended by Tong et al. [17]. The expressions levels of seven genes are low (*Unknown 3* and *Unknown 7*) and intermediate (*RP II, PRT 6, TUA 3, ATPase P* and *GTB 1*). None of the genes in this set had high expression levels. They were discarded early from among the 21 preselected genes because of their high Cq levels under hypoxia stress using NormFinder complement (Table 4–5). Therefore, as [34] described, this approach can also validate low and intermediate expression housekeeping genes, in contrast to more “traditional” housekeeping genes whose expression is significantly higher than that of the genes of interest.

Roots are composed of different cellular zones. Three zones have been identified to date along the longitudinal axis of the primary Arabidopsis root: the root apical meristematic zone (RAM) with two domains [the proliferative (PD) and the transition domains (TD)], the elongation zone (EZ), and the maturation zone (MZ) [35,36]. These different zones need to be considered when determining suitable reference genes, especially in developmental genetic studies of root morphogenesis. These results will facilitate new genic expression studies of *Prunus sp.* that can improve our understanding of the molecular mechanisms of plants under abiotic or other stress.

This study validated housekeeping *Prunus sp*. genes for normalizing gene expression analysis with RT-qPCR, which can be used in new gene expression studies. Our results suggest that different combinations of suitable housekeeping genes should be selected for normalization according to the genotypes, tissue or treatments to be evaluated.

## Acknowledgments

This work was funded by FONDECYT Project 1161377. AB was supported by FONDECYT Project 11150551. RA acknowledge to CEAF_R08I1001. Plant materials were kindly provided by Agromillora Sur S.A.

